# A Novel Conjugation System in AMR-associated pELF-type Linear Plasmids of *Enterococcus faecium*

**DOI:** 10.64898/2026.01.27.701956

**Authors:** Jun Kurushima, Natsuko Ota, Yuka Yoshii, Naoko Tomie, Haruyoshi Tomita

## Abstract

The pELF-type linear plasmid is a unique transconjugative plasmid that disseminates various antimicrobial resistance (AMR) genes (e.g., vancomycin resistance in vancomycin-resistant *Enterococcus faecium*, [VRE]), leading to hospital outbreaks worldwide. Despite evidence of its role in AMR gene expansion, the molecular mechanisms underlying conjugative transfer of pELF-type plasmids have been poorly understood due to their unique linear structure and the lack of convenient genetic engineering tools adapted to *E. faecium* strain. Herein, we focused on a putative transconjugation-associated gene cluster, named the *tra*_pELF_ cluster, encoded on pELF2, a *vanA*-type vancomycin resistance gene-harboring plasmid. RNA-seq analysis of the pELF2 plasmid identified a putative operon containing an *ftsK*-like ATPase-encoding gene and an additional 10 genes as candidates for conjugative transfer of pELF-type linear plasmids. We constructed isogenic deletion mutants for these 10 genes and revealed that two genes, designated *traC* and *traD*, are essential for the conjugative transfer of pELF2 to a recipient strain. Inducible overexpression of the *tra* genes *in trans* from an ectopic shuttle vector confirmed the essentiality of *traC* and *traD* for conjugative transfer. Promoter assay using a luciferase reporter plasmid revealed that the upstream region of the putative *tra* operon possesses functional promoter activity characterized by constitutive strong transcriptional expression. Furthermore, we observed membrane localization for a fluorescent protein fusion of TraC, but not for that of TraD. Based on these findings, we propose a working model for the unique molecular machinery for the transconjugation of pELF-type linear plasmids.

**Author Summary:** Enterococci are among the opportunistic pathogens of greatest global concern due to their multidrug resistance, and it is well established that various plasmids mediate horizontal transfer of resistance genes in this species, converting susceptible strains into resistant ones. While most enterococcal plasmids have a circular structure, a linear pELF-type plasmid was recently discovered. Since then, pELF-type plasmids have been reported worldwide as important vehicles for antimicrobial resistance genes, but their mechanism of transfer has remained unknown. In this study, we have, for the first time, successfully identified the genes required for the conjugative transfer of pELF-type plasmids. We confirmed that these genes are absent from known conjugative plasmids, yet are highly conserved among pELF-type plasmid sequences deposited in public databases. Taken together, these findings suggest that pELF-type plasmids possess a unique transfer mechanism that reflects their distinctive molecular architecture.

## Introduction

*Enterococcus faecium* is a Gram-positive, coccoid bacterial species that commonly inhabits the gut microbiota in healthy humans and is also widely distributed across natural environments [1]. However, *E. faecium* is an opportunistic pathogen, and vancomycin-resistant *E. faecium* (VRE) has emerged as one of the most concerning antimicrobial-resistant (AMR) bacterial pathogens [2]. Among the related bacteria, *Enterococcus* species are known to harbor large plasmids, which drive rapid genetic evolution by enabling the acquisition of new genes, such as those conferring antimicrobial resistance [3]. In *E. faecium*, horizontal gene transfer of AMR genes (ARGs) is mediated by conjugative circular plasmids with various *rep* types, including pMG1-like plasmid (e.g., pMG1, pHTβ, and pZB18) or pRepA_N-group plasmid (e.g., pRUM-type plasmids) [4–22].

Phylogenomic studies revealed that *E. faecium* is divided into clades A1, A2, and B [23]. Clade A1 is hospital-associated with a high risk for enterococcal infection and contains the clonal complex 17 (CC17) lineage which is recognized as the highest risk lineage for human health according to the conventional multi-locus sequence typing (MLST) method [24]. Clade A2 is phylogenetically close to clade A1; however, it is primarily associated with animal hosts rather than human infection. Clade B *E. faecium* is genetically distinct from the Clade A lineages**—**having been recently re-designated as the species *Enterococcus lactis***—**and is a major symbiont bacterial species in the human gut [25]. Notably, despite these differences in the clinical risk, all *E. faecium* clades and *E. lactis* share conjugative plasmid types that harbor the ARGs [26,27]. These findings highlight that horizontal transfer via conjugative plasmids plays a key role in the dissemination of the ARGs among *E. faecium* and *E. lactis* strains over various ecological sectors, including human, animals and environmental resources [28].

The pELF-type plasmid was recently discovered and is a large (generally > 100 kb) linear DNA molecule that allows multiple genetic elements to be integrated [29]. Interestingly, pELF1 is the first reported pELF-type plasmid, isolated from a hospitalized patient in Japan and harboring both *vanA*- and *vanM*-type vancomycin resistance gene cluster [7]. pELF-type linear plasmids have been frequently isolated in association with multiple ARGs, including those conferring glycopeptides and linezolid resistance [30–37]. Besides antimicrobial resistance, the pELF-type plasmid has also been reported to harbor metabolic gene sets that confer a competitive survival advantage in the human intestine [38]. The pELF-type plasmids have been reported to be highly conjugative among *Enterococcus* species and stably maintained under laboratory conditions, although *E. faecium* and *E. lactis* appear to be their natural hosts [29]. Notably, pELF-type linear plasmids lack the conventional conjugation type IV secretion system (T4SS) gene sets that were characterized in circular conjugative plasmids [39] or even the linear conjugative plasmids of *Streptomyces* [40,41]. In this study, we analyzed the genetic functions of the pELF-type linear conjugative plasmid family and identified novel transconjugation genes that are essential for the conjugative transfer of the pELF plasmid.

## Results

### Transcriptome analysis identified a putative conjugation operon in the pELF-type linear plasmid

Strain KUHS13 is a clade A1-lineage *E. faecium* isolate that harbors the pELF2 plasmid (<u>AP022343.1</u>), which is a pELF-type linear plasmid carrying Tn*1546* encoding the *vanA*-type vancomycin resistance operon [42]. To analyze the transcriptional profile of genes encoded on the pELF2 plasmid, we isolated total RNA from KUHS13, a human-adapted lineage *E. faecium* strain harboring pELF2 during the exponential (4 h) or stationary (8 h) phases in culture in brain heart infusion (BHI) broth (Supplementary Fig S1). Furthermore, to investigate the effect of the co-presence of the recipient cell on transconjugation, total RNA was isolated from co-cultures with the plasmid-null strain BM4105RF, a laboratory model *E. faecium* strain exhibiting rifampicin- and fusidic acid-resistance. Differential expression analysis reported only a limited change in transcriptional profiles among these conditions (Fig 1A and B).

**Fig 1.**
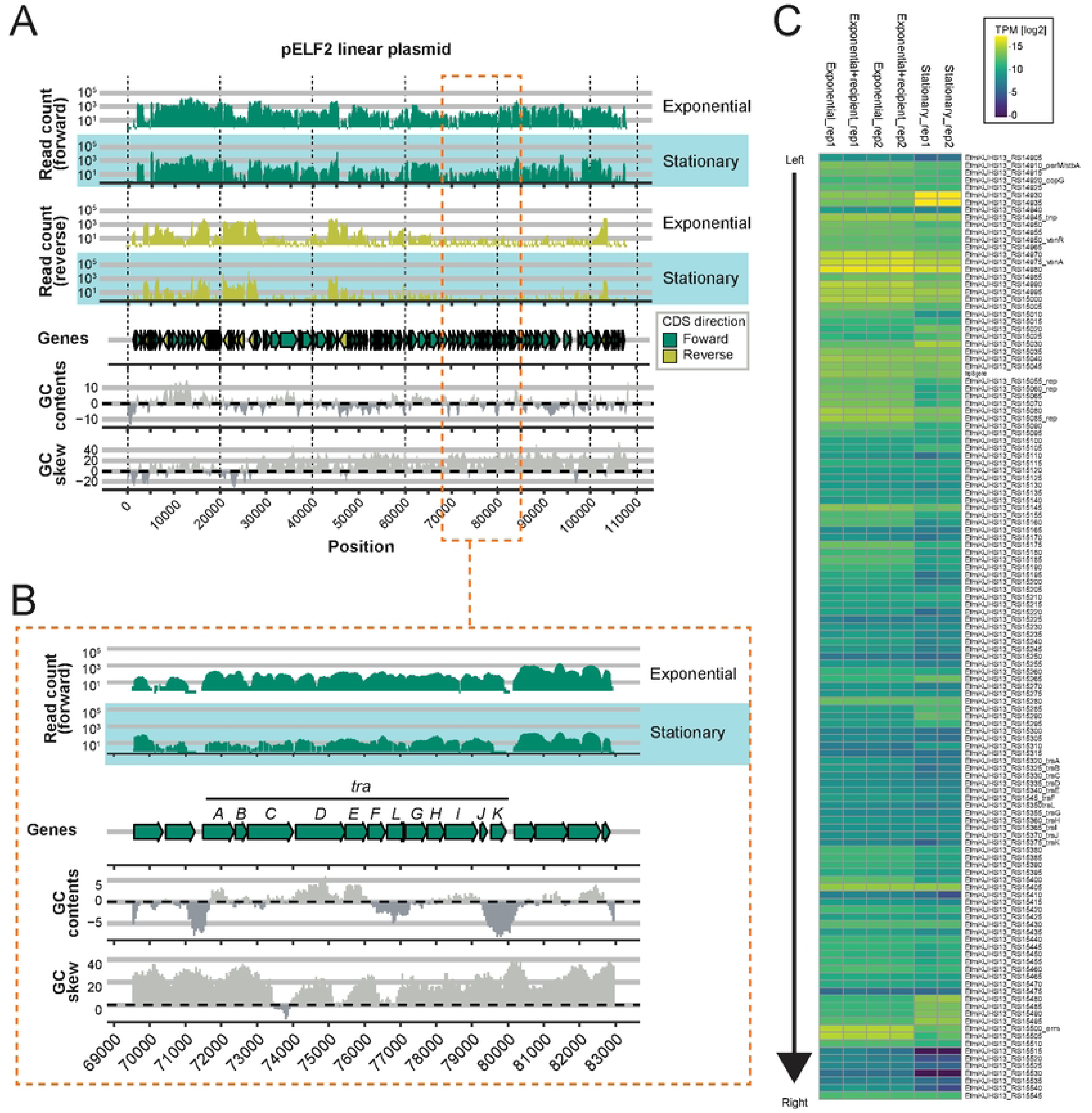
Transcriptional profile of the pELF2 linear plasmid. (A) Schematic presentation of the RNA-seq coverage plot on pELF2 plasmid. Mapping data for the forward or reverse strand, as well as annotated coding sequences (CDSs) located on the forward or reverse strands, are indicated in green and yellow, respectively. Bottom panels display GC contents and GC skew across the pELF2 nucleotide sequence. (B) Enlarged view of panel A, focusing on the *tra*_pELF_ genes region to provide greater detail of its features. (C) Gene expression matrix for strain KUHS13 during the exponential growth phase (with or without recipient cell BM4105RF) and the stationary growth phase.

RS15335 in pELF2 encodes a homolog of the FtsK-type ATPase, also known as a type IV coupling protein (T4CP). RNA-seq read coverage suggested that RS15335 is transcribed as part of an operon-like structure containing 11 putative coding sequences (CDSs), designated *traA–L* (Table 1). We designated this putative operon transconjugation (*tra*_pELF_) operon as a candidate genetic element involved in the conjugative transfer of pELF-type plasmids (Fig 1C). In contrast to the T4CP homolog (*traD*), the remaining CDSs showed no similarity to conventional plasmid conjugation genes. Transmembrane domains were detected in the gene products of *traC*, *traG*, *traI*, and *traK*. The *traA* gene product, TraA contains a non-specific endonuclease domain. Both TraG and TraL contain a rhomboid-family intramembrane serine protease domain. Notably, *traL*, which is encoded immediately downstream of *traG* with a slight sequence overlap, was not previously annotated in the publicly available data (<u>AP022343.1</u>) as a CDS. The remaining gene products (TraB, TraC, TraE, TraF, TraH, TraI, TraJ, and TraK) lacked known functional domains.

**Table 1.** Genes encoded in *tra*_pELF_ operon.

| Gene | Start<sup>1</sup> | End<sup>1</sup> | length (nt) | length (a.a.) | Functional domain | TM domain<sup>2</sup> | Role on conjugation<sup>3</sup> |
| --- | --- | --- | --- | --- | --- | --- | --- |
| *traA* | 71'490 | 72'359 | 870 | 289 | DNA/RNA nonspecific endonuclease | None | Not tested |
| *traB* | 72'379 | 72'726 | 348 | 115 |  | None | Not tested |
| *traC* | 72'739 | 74'001 | 1'263 | 420 |  | 2 | Essential |
| *traD* | 74'081 | 75'457 | 1'377 | 458 | FtsK-type ATPase | None | Essential |
| *traE* | 75'454 | 76'083 | 630 | 209 |  | None | Not detected |
| *traF* | 76'102 | 76'581 | 480 | 159 |  | None | Not detected |
| *traL* | 76'623 | 77'162 | 540 | 179 | Rhomboïd family serine protease | None | Not tested |
| *traG* | 77'140 | 77'739 | 600 | 199 | Rhomboïd family serine protease | 6 | Not detected |
| *traH* | 77'755 | 78'231 | 477 | 158 |  | None | Positive |
| *traI* | 78'248 | 79'150 | 903 | 300 |  | 2 | Not detected |
| *traJ* | 79'223 | 79'417 | 195 | 64 |  | None | Not tested |
| *traK* | 79'522 | 79'950 | 429 | 142 |  | 1 | Not tested |

<sup>1</sup> Positions are indicated from nucleotide sequence of pELF2 plasmid
<sup>2</sup> Trans membrane (TM) domains were detected with TMHMM v2.0 (https://services.healthtech.dtu.dk/services/TMHMM-2.0/)
<sup>3</sup> Roles on conjugation refer to the results in Figure 2

### Essentiality of the *tra* genes for the conjugative transfer of pELF2

Although the *tra* operon contains 11 CDSs, the encoded proteins exhibit no similarity to known plasmid-conjugation proteins. To investigate the essentiality of the *tra* genes, we constructed isogenic deletion mutants for every gene within the *tra* operon. To construct an in-frame deletion mutants without selection markers, we developed a suicide plasmid (pMGJK72) that enables an improved targeted genetic recombination system optimized for *E. faecium* (see Materials and Methods for details; Supplementary Fig S2). Deletion mutants for *traA*, *traB*, *traG*, *traJ*, and *traL* could not be generated. The obtained deletion mutant collection (derived from the parent strain KUHS13) harboring pELF2 as the original host was evaluated as donor strains in conjugative transfer experiments under both filter and broth conditions, using BM4105RF as the recipient strain (Fig 2A). These gene deletions did not impact bacterial growth (Supplementary Fig S3). The wild-type parent strain, KUHS13, exhibited transfer efficiencies of 1.0 × 10^-3^ and 1.0 × 10^-6^ in filter and broth mating, respectively. Deletion of *traC* (Δ*traC*) and *traD* (Δ*traD*) completely abolished conjugative transfer. Deletion of *traH* (Δ*traH*) also resulted in a 10-fold reduction in transfer efficiency compared to that of the wild-type under both filter and broth mating conditions. The remaining mutants (Δ*traE, ΔtraF, ΔtraI*, and Δ*traK*) showed no significant differences compared to the wild-type strain (KUHS13).

**Fig 2.**
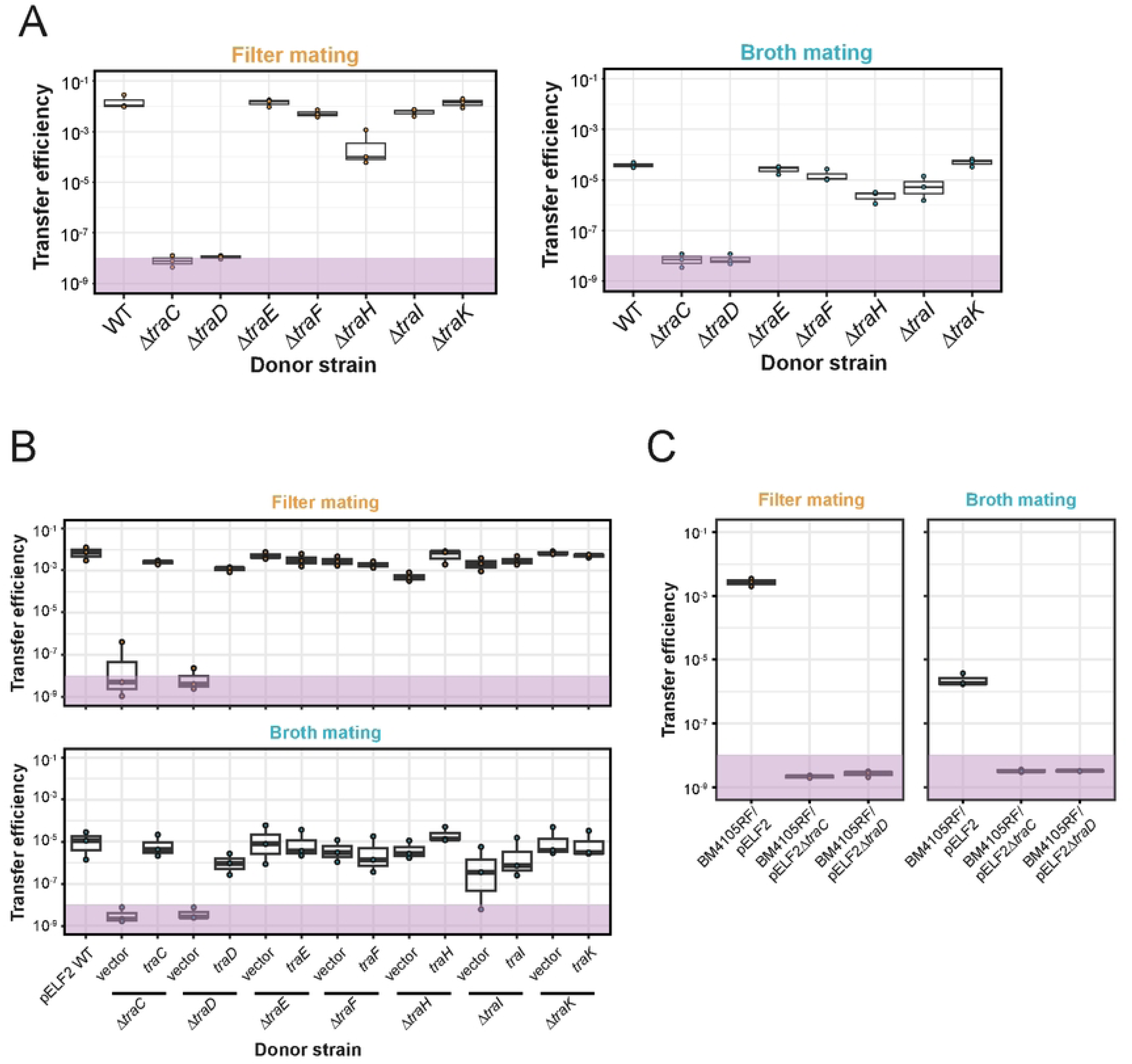
Effect of the *tra* gene deletions on the conjugative transfer of pELF2. (A) Conjugation efficiency of pELF2 plasmid derivatives with individual in-frame deletions. Donor strain KUHS13 harboring pELF2 derivatives and recipient strain BM4105RF were mixed and incubated under filter (left panel) or broth (right panel) conditions. (B) Conjugation efficiency of the pELF2 plasmid with individual in-frame deletions harboring cognate complementation vectors. Donor strain KUHS13 harboring pELF2 derivatives and recipient strain BM4105RF were mixed and incubated under filter (top panel) or broth (bottom panel) conditions. IPTG induction was performed during pre-incubation and co-cultivation for conjugation. (C) Secondary conjugation efficiency of BM4105RF donors harboring pELF2 derivatives. Donor strain BM4105RF harboring pELF2 derivatives and recipient strain BM4105SS were mixed and incubated under filter (left panel) or broth (right panel) conditions. For values below the detection limit, the limit point is plotted as a proxy (purple area). Data represent the means and error bars from three independent replicates.

To confirm these phenotypes, we developed an IPTG-inducible ectopic expression vector (pMGJK53) optimized for *E. faecium* (Supplementary Figs S4 and S5) and performed complementation assays for each deletion mutant (Fig 2B). No differences in conjugation efficiency were detected between wild-type KUHS13 with and without the control pMGJK53 vector (pMGJK53), indicating that neither the introduction of the pMGJK53 vector nor the presence of IPTG interfered with conjugative transfer efficiency. Complementation using either pMGJK53::*traC* or pMGJK53::*traD* restored the impaired conjugation phenotype in the *ΔtraC* and *ΔtraD* strains, respectively. Notably, the resulting transconjugants BM4105RF harboring pELF2*ΔtraC* or pELF2*ΔtraD* were incapable of secondary conjugation to the BM4105SS strain (Fig 2C). This result ruled out the possibility that unintended mutations or integration events in pELF*ΔtraC* or *ΔtraD* were responsible for the restoration of the conjugation capacity. The decreased transconjugation efficiency in the *ΔtraH* mutant was also restored to wild-type levels via the introduction of pMGJK53::*traH*. Furthermore, ectopic overexpression of the individual *tra* gene in the wild-type KUHS13 background yielded no significant differences compared to wild-type KUHS13 lacking an expression vector or harboring the control pMGJK53::*gfp* vector (Fig 3).

**Fig 3.**
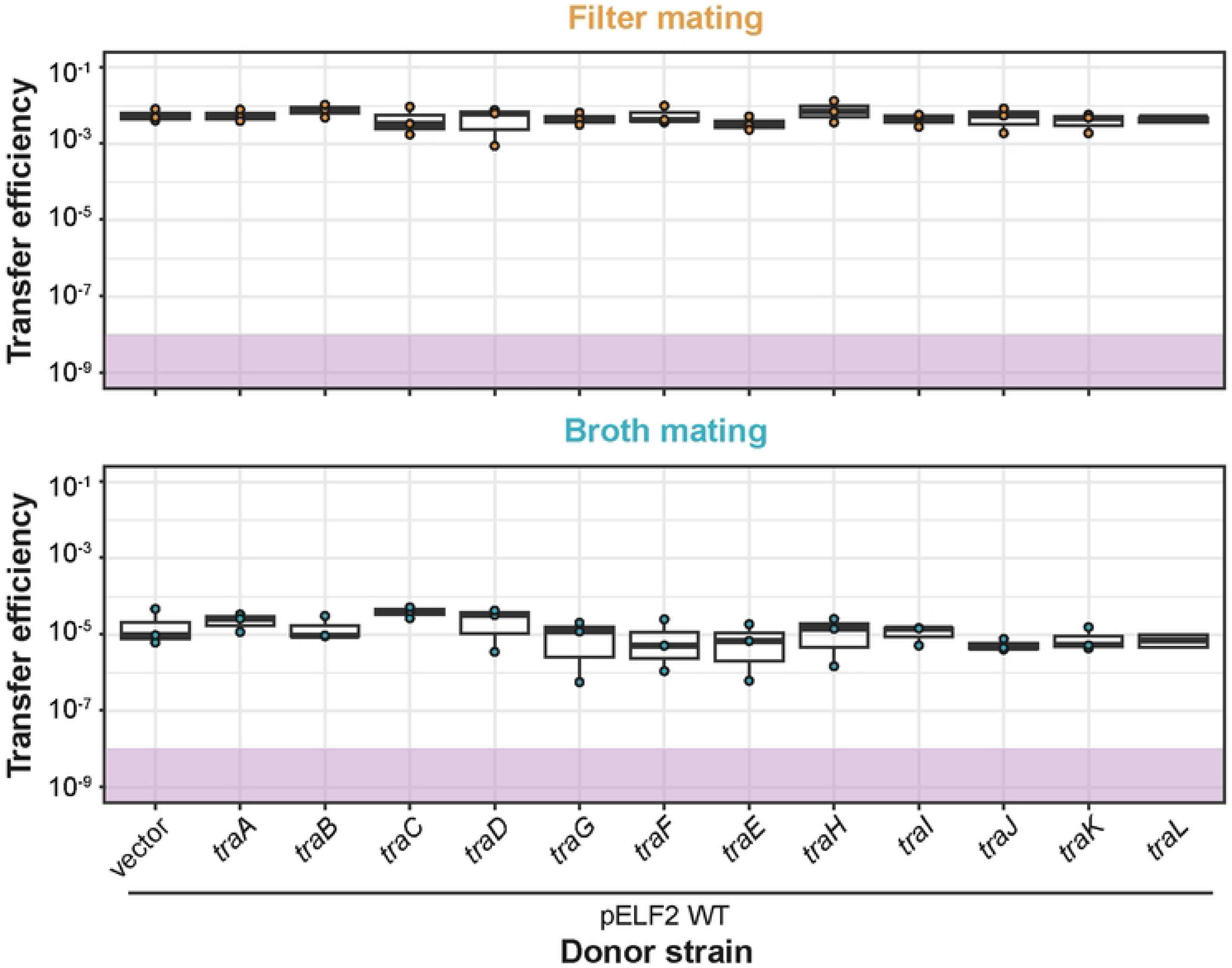
Effect of the *tra* genes overexpression on the conjugative transfer of pELF2. Conjugation efficiency of the pELF2 plasmid during the overexpression of individual *tra* genes. Donor strain KUHS13 (harboring the overexpression vector) and recipient strain BM4105RF were mixed and incubated under filter (top panel) or broth (bottom panel) conditions. Values below the detection limit, the limit point is plotted as a proxy (purple area). Data represent the means and error bars from three independent replicates.

Collectively, these data demonstrate that both *traC* and *traD* are essential for the conjugative transfer of pELF2, and that *traH* positively impacts conjugative transfer. No involvement of *traE*, *traF*, *traI*, or *traK* genes in the conjugative transfer of pELF2 was detected, at least under the tested conditions.

### Identification of a functional promoter upstream of the *tra* gene

Pheromone-responsive conjugative plasmids in *E. faecalis* are the most widely studied type of enterococcal plasmids; their conjugation is transcriptionally induced in response to quorum-sensing peptide pheromones produced by recipient cells. The operon-like expression pattern observed in the RNA-seq data suggests a transcriptional start site at approximately position 71,468 bp upstream of the *traA* gene (Figs 1 and 4A). To assess putative promoter activity for the *tra* operon, fragments of the putative promoter region were cloned into the pMGJK55 plasmid, a luciferase reporter vector designed for promoter assays (Supplementary Fig S6). Fragments P*tra*1, P*tra*2, and P*tra*3 were generated by amplifying the regions from positions 71,256, 71,358 and 71,406 to 71,457 of the pELF2 plasmid, respectively (Fig 4A). Wild-type KUHS13 was transformed with the reporter constructs, and bioluminescence activity was monitored during bacterial growth using a plate reader (Fig 4B). The promoterless negative control construct produced no detectable bioluminescence signal. All the tested promoter-fragment constructs exhibited higher signals than that of the positive control (P*lac* promoter in the absence of the *lacI* repressor gene) (Fig 4B). To determine whether individual *tra* genes regulate the promoter activity, the P*tra*1-cloned construct was introduced into the *traC*, *traD*, *traE*, *traF*, *traG*, *traH*, *traI*, and *traK* deletion mutants (Fig 4C). None of these mutant backgrounds affected the P*tra*1-driven expression signal, suggesting that these *tra* genes do not function as regulators of the *tra* operon expression.

**Fig 4.**
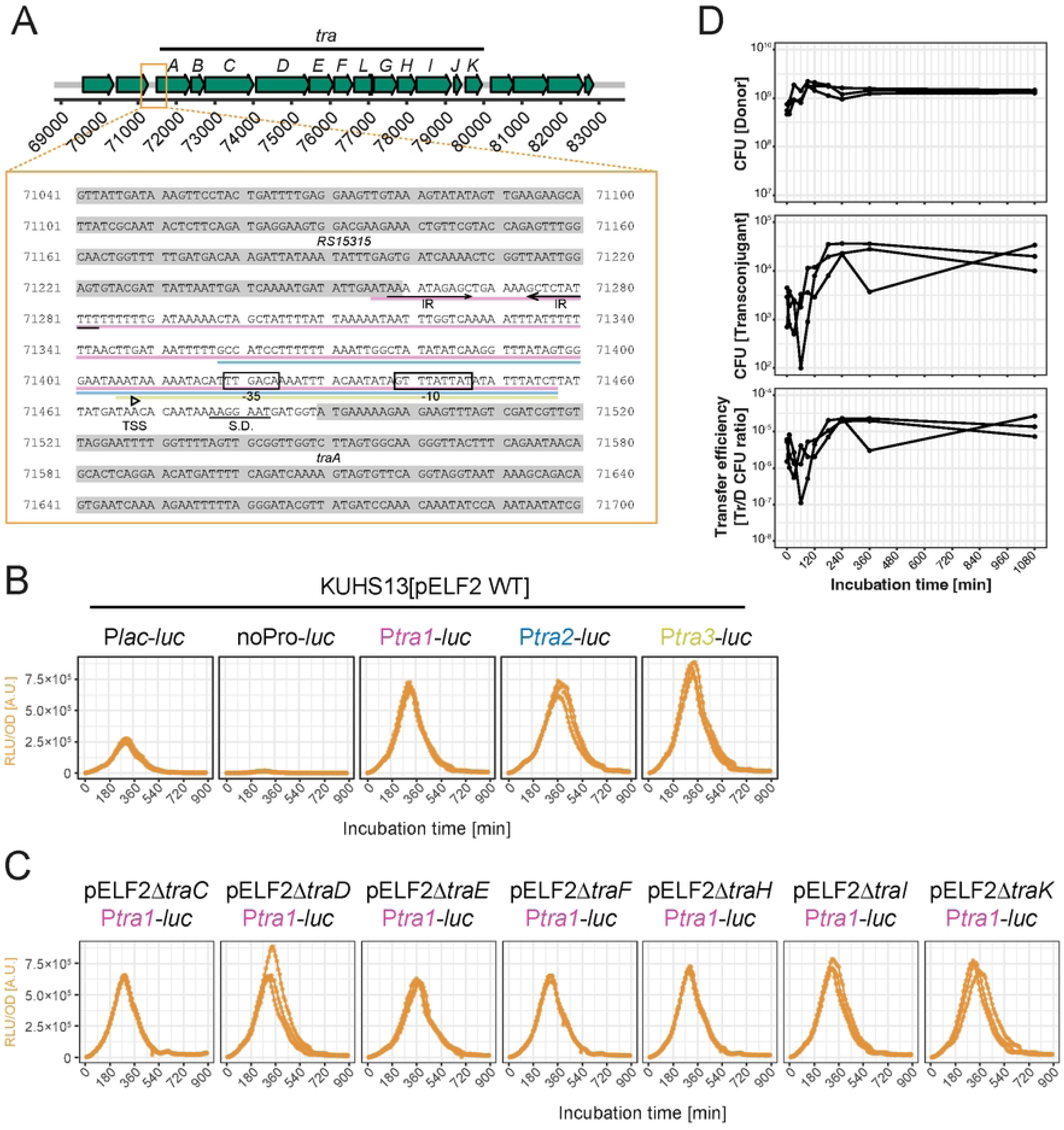
Promoter activity of the upstream region of the putative *tra*_pELF_ operon. (A) Nucleotide sequence of upstream region of *traA* gene. The putative transcription start site (TSS) was determined from the RNA-seq coverage. Cloned promoter regions (P1, P2, and P3) are highlighted with pink, blue and yellow underlines, respectively. (B) Kinetics of promoter activation. OD-normalized bioluminescence activity of the wild-type KUHS13 harboring pMGJK55::P*tra*1, pMGJK55::P*tra*2, or pMGJK55::P*tra*3 in the presence of 10 µg/mL D-luciferin. Plots represent values of three replicates. (C) Kinetics of promoter activation in individual *tra* gene mutant backgrounds. OD-normalized bioluminescence activity of the KUHS13 derivatives harboring pMGJK55::P*tra*1 in the presence of 10 µg/mL D-luciferin. Plots represent values of three replicates. (D) Kinetics of pELF2 transfer. Broth mating experiment was performed as described above, with sampling for plating at each time point. Plots represent values of three replicates.

### Subcellular localization of fluorescently labeled TraC and TraD proteins

The newly identified conjugative genes, *traC* and *traD*, appear to play no role in the *tra* operon transcription. Since TraD is a homolog of the FtsK-type T4CP motor, it likely functions as a part of a plasmid DNA translocation system. The protein complexes required for plasmid conjugative transfer must assemble across the cellular membrane and cell wall to reach the recipient cell surface. Amino acid sequence analysis indicated that transmembrane domains are present in TraC but not in TraD (Fig 5A). To analyze the subcellular localization of TraC and TraD, each gene was cloned into pMGJK53 to express a C-terminal fusion with mScarlet-I. The mScarlet-I fusion constructs were introduced into wild-type KUHS13 as well as the *traC-* or *traD*-deficient mutants.

**Fig 5.**
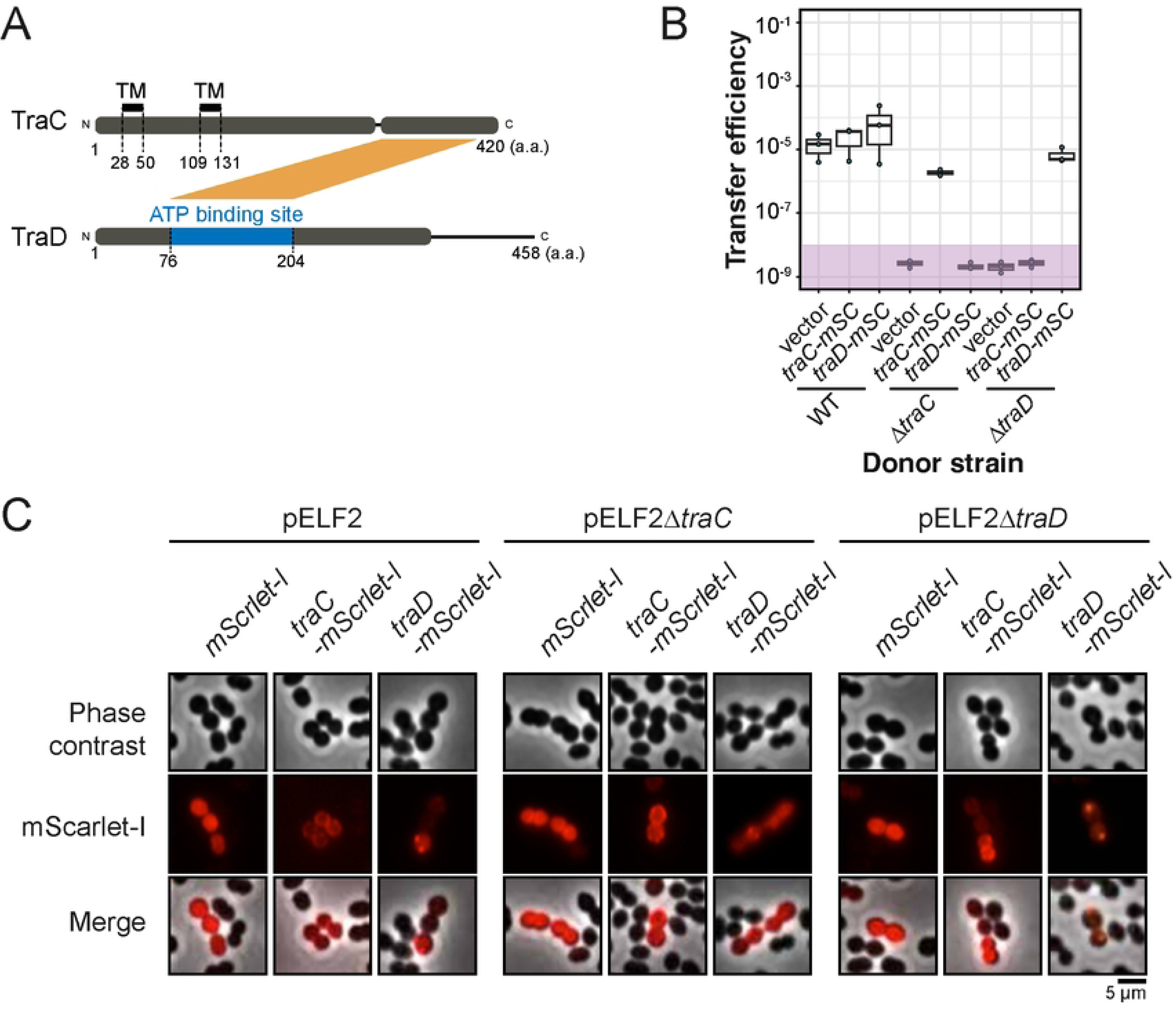
Subcellular localization of the mScarlet-I-fused TraC and TraD. (A) Molecular structures of TraC (420 amino acids, bottom) and TraD (458 amino acids, top) proteins with conserved functional domains. TraD contains an ATP-binding domain, whereas TraC contains two transmembrane domains. (B) Conjugation efficiency of pELF2 in-frame deletion mutants harboring mScarlet-I fusion protein expressing vectors. Donor strain KUHS13 harboring pELF2 derivatives and recipient strain BM4105RF were mixed and incubated under broth conditions. IPTG induction was performed during pre-incubation and co-cultivation for conjugation. Values below the detection limit, the limit point is plotted as a proxy (purple area). Data represent means and error bars from three independent replicates. (C) Fluorescent microscopy images of *E. faecium* KUHS13 expressing TraC or TraD mScarlet-I fusions in the wild-type (left panels), *traC* mutant (middle panels) or *traD* mutant (right panels). Scale bar: 5 µm.

Notably, the mScarlet-I fusions did not abolish conjugative transfer (Fig 5B). After incubation in the presence of 0.1 mM IPTG, the cells were collected and analyzed via fluorescence microscopy (Fig 5C). Expression of the mScarlet-I protein alone showed a diffuse signal throughout the cytosol. In all tested genetic backgrounds, the TraC signal localized to the cytoplasmic membrane. Conversely, the TraD signal formed discrete foci in the wild-type and *ΔtraD* background. In the *ΔtraC* background, the TraD signal became diffuse. These observations suggest that TraC localizes to the cytoplasmic membrane and that TraD assembles into discrete foci in a *traC-*dependent manner.

### Phylogenetic conservation of the *tra* operon among pELF-like plasmids

To investigate whether the *tra* genes are phylogenetically conserved among the pELF-type plasmid family, 2,060 replicon contigs were retrieved from 675 *E. faecium* strain genomes deposited in the RefSeq microbial genome database [43]. Of these, 57 contigs contained the 500-bp left-hand hairpin nucleotide sequence, which is a hallmark of pELF-type plasmids. In the pan-genomic analysis of these 57 high-quality pELF-like plasmid sequences, 46 conserved genes, including those involved in replication and partitioning, were identified (Fig 6). The nine genes (*traB*, *traC*, *traD*, *traE*, *traG*, *traH*, *traI*, *traJ* and *traL*) of the *tra*_pELF_ cluster are conserved. The *traF* gene was highly conserved, appearing in 50 of the tested pELF-like plasmid sequences (89.3%). Conversely, the *traA* and *traK* genes, located at the head and tail of the *tra*_pELF_ operon, were present in the 42 and 17 sequences, respectively. Plasmids NZ_CP019973.1, NZ_CP019995.1, and NZ_CP058893.1 completely lack the *tra_pELF_* operon as well as a large portion of the right-hand region (Fig 6).

**Fig 6.**
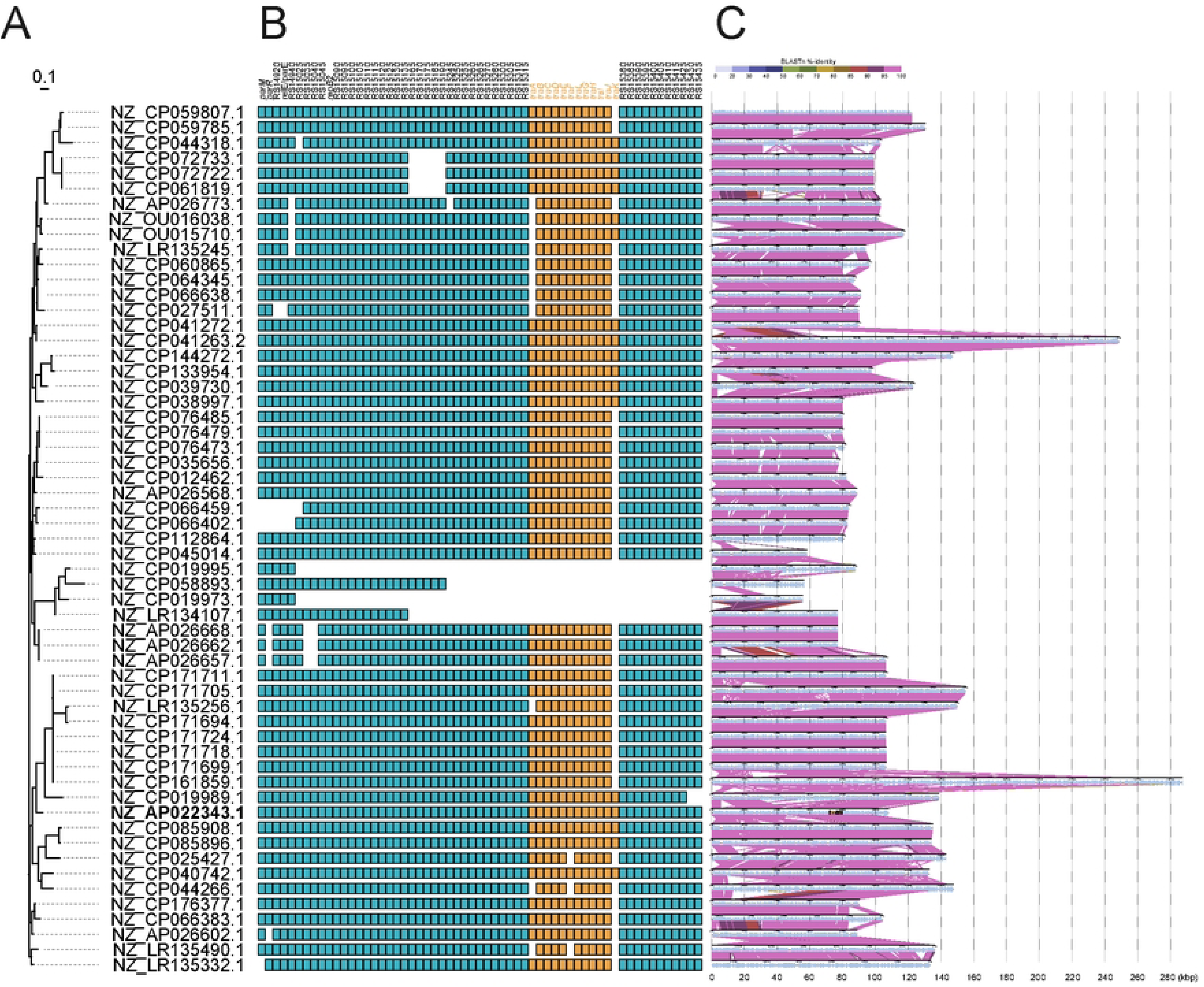
Phylogenetic analysis of the *tra* genes among pELF-like plasmid sequences. (A) Phylogenetic tree of the left-hand hairpin structure-positive plasmids retrieved from the NCBI RefSeq database. pELF2 (NZ_AP022343.1) is indicated in bold. (B) Presence of the 11 *tra* genes (orange) and other conserved genes (blue). CDS annotation was performed using Roary. (C) Synteny block plots were generated with DiGAlign. Similarity between nucleotide sequences was calculated via BLASTn.

## Discussion

In this study, we analyzed RNA-seq data and identified a putative operon likely related to the conjugative transfer of pELF2. This operon, named the pELF-type linear plasmid-related *tra* operon (*tra*_pELF_ hereafter). It encodes 11 CDSs, *traA–L*, including a VirD/FtsK-type ATPase motor homolog, *traD*_pELF_. By constructing isogenic gene deletion derivatives of the pELF2 plasmid, we confirmed that *traD*_pELF_ is essential for the conjugative transfer of pELF2. Additionally, the hypothetical protein-encoding gene *traC*_pELF_ was revealed as a novel essential conjugative gene of pELF2. Furthermore, deletion of *traH*_pELF_ resulted in a partial decrease in conjugation efficiency, suggesting that *traH* is not essential but has an unknown positive role in the transconjugation of pELF2. Meanwhile, deletion mutants of *traE*_pELF_, *traF*_pELF_, *traI*_pELF_, and *traK*_pELF_ were obtained but exhibited no effect on transconjugation, suggesting that these genes do not contribute, at least in this experimental setting, to conjugative transfer. The roles of *traA*_pELF_, *traB*_pELF_, *traG*_pELF_, *traJ*_pELF_, and *traL*_pELF_ in the conjugative transfer of pELF2 could not be assessed in this study and remain subject to future research because deletion mutants of these genes could not be constructed.

Promoter analysis of the upstream region of *traA*_pELF_ suggested that the transcription of the putative *tra*_pELF_ operon is coordinated by at least a 100-bp region from the speculated transcriptional start site (Fig 4). Meanwhile, no effect from the deletion of the *traC*_pELF_, *traD*_pELF_, *traE*_pELF_, *traF*_pELF_, *traH*_pELF_, *traI*_pELF_, or *traK*_pELF_ was observed, and the transcription appeared to be constitutively induced throughout bacterial growth regardless of the presence of recipient cells (Figs 1 and 4). The recipient-dependent induction of conjugative gene expression has been well-studied in pheromone-responsive conjugative plasmids in *E. faecalis* [44]. In this system, a peptide composed of eight amino acids is chromosomally produced from *E. faecalis* and is recognized by a cognate pheromone-responsive plasmid to induce transcription of the conjugative operon [45,46]. On the other hand, induction modules stimulated by external signals have not been identified in *E. faecium* conjugative plasmids (e.g., pMG1-like plasmids) [47]. However, it is quite possible that such a mechanism will be discovered as research on *E. faecium* plasmids progresses. Furthermore, more direct evidence, such as Cappable-Seq, 5′-RACE assay, or Northern hybridization, is necessary to establish transcriptional operon structure for the *tra*_pELF_ gene cluster.

In general, bacterial conjugative plasmid transfer is carried out by a relatively large protein complex that localizes to the donor cell membrane to form a molecular channel for passing the plasmid DNA substrate to the recipient cell [48]. The classical conjugative transfer model for circular plasmids, including enterococcal plasmids, is that a plasmid-encoded nickase (also known as a relaxase) generates single-stranded DNA at an internal site of the substrate plasmid and subsequently loads it into the conjugative type IV secretion system (T4SS) apparatus for transfer to the recipient cells [49]. However, since this model relies on the rolling-circle replication (RCR) mechanism, it is not applicable to the transfer of linear plasmid DNA. In contrast with conventional circular plasmids, knowledge of the conjugation mechanisms of linear plasmids has been limited; however, genetic studies of SLP2 from *Streptomyces* provide insights into linear plasmid transfer [50]. The SLP2 plasmid that plays a role in horizontal transfer of secondary metabolic genes, such as those for antibiotic production in *Streptomyces*, is one of the most-studied conjugative linear plasmids. Conjugation of SLP2 is not mediated through conversion into a single strand; rather, it is thought to occur via the first-end model, in which the linear plasmid molecule, with its ends bound by protective proteins (telomere or terminal proteins), is transferred to the recipient in a double-stranded form through the conjugative transfer apparatus starting from one end [51]. Since no nickase homolog is present in the pELF-type plasmids, a first-end model similar to that of SLP2, rather than RCR, appears to be the most plausible explanation. Notably, the components of the conjugative transfer apparatus themselves exhibit limited similarity, and the terminal structures of the linear plasmid DNA differ slightly between pELF and SLP2. Therefore, further studies are needed to clarify the physical state of the pELF molecules during conjugative transfer.

The studies of the SLP2 conjugative linear plasmid reported that a FtsK/VirD4-type ATPase motor is the only essential factor for its conjugative transfer. The FtsK/VirD4 homolog from SLP2 shares only a limited portion of the ATPase domain with TraD_pELF_; however, the genetic contexts around the genes encoding these FtsK-like translocators show no similarity. Instead, protein structure modeling using AlphaFold3 (https://alphafold.ebi.ac.uk/) algorithm predicted a stable interaction between the C-terminal portion (approximately 300–400 a.a.) of TraC_pELF_ and the ATPase domain (approximately 30–320 a.a.) of TraD_pELF_ (Fig 5A) [52], suggesting that TraD_pELF_ is anchored to the cytoplasmic membrane via interaction with TraC (Fig 7D). This hypothesis may be supported by the fluorescent microscopic images in Fig. 5, where TraD_pELF_ localization appears altered in the *traC* deletion background. Unfortunately, in the current pMGJK53 expression system developed in this study, fluorescence signal intensity is heterogeneous across cells, and only a limited number of cells exhibit clearly observable localization. This makes statistical quantitative analysis of localization patterns difficult. Future improvements to the expression system will likely enable detailed examination of the intracellular localization of each protein component.

**Fig 7.**
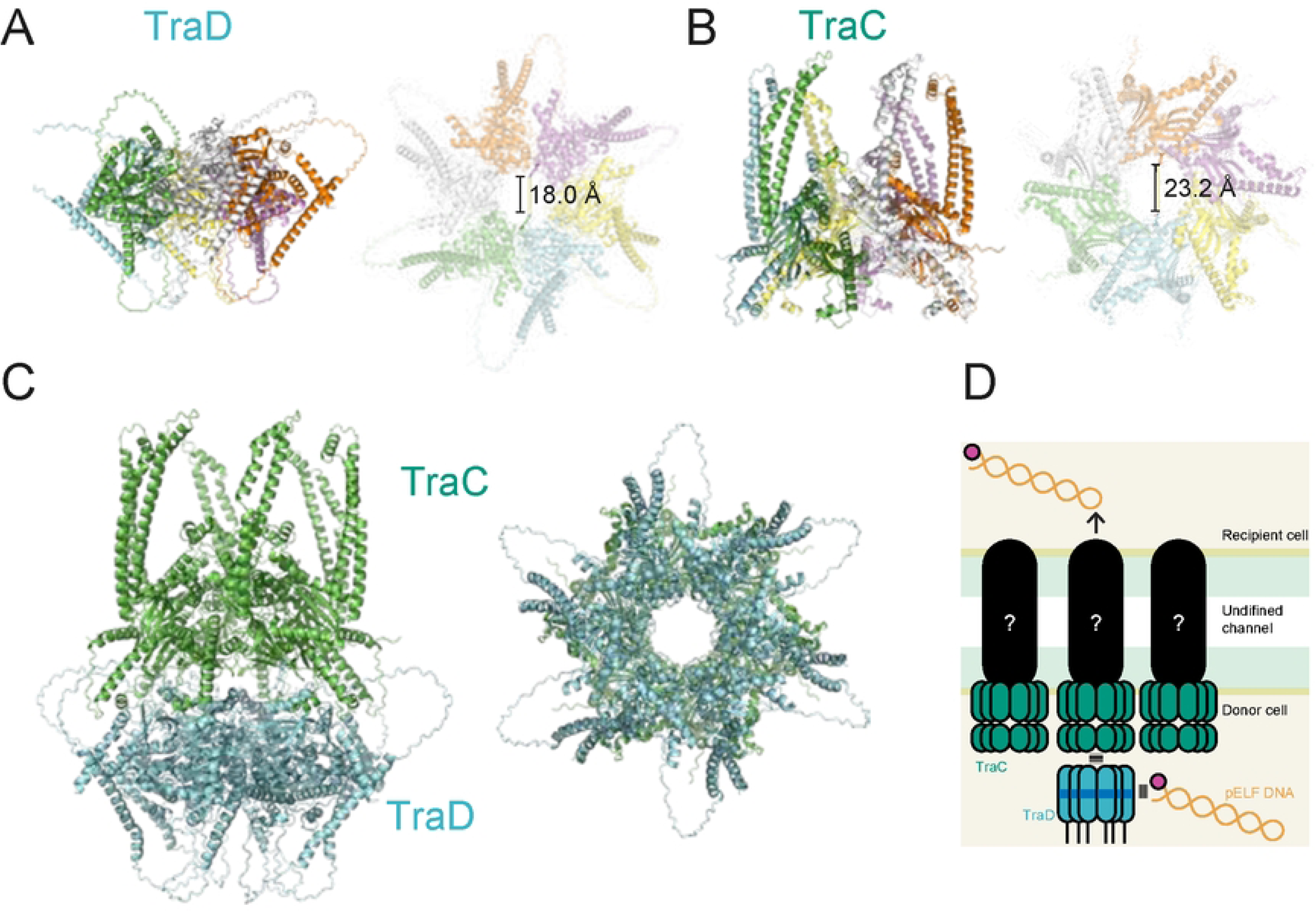
Proposed working model for the conjugation machinery of pELF-type plasmid. Protein structure modeling of TraD (A) or TraC (B) homohexamer generated from six copies of each protein sequence using AlphaFold3. TraC and TraD sequences are from pELF2 (NZ_AP022343.1). (C) TraC and TraD each form stable homohexamers. (D) TraD possesses a DNA-binding ATPase domain, which can capture plasmid DNA for transfer; however, it cannot localize to the cell membrane independently. Therefore, through interaction with the membrane-associated TraC, TraD is thought to load plasmid DNA into a putative, yet-unidentified secretion apparatus within the cell membrane. The purple circle at the end of pELF plasmid DNA molecule indicates putative terminal protein.

Conventional bacterial T4SSs are divided into two major types, defined by their structural basis [53]. Type A is a conjugation T4SS represented by *E. coli* F plasmids R388 and pKM101. Type B T4SS is directly relevant to virulence and represented by the *Legionella* Dot/Icm system, which functions as a virulence effector protein exporter to host cells. Although T4SSs are found in many bacterial species and share conserved structural components such as ATPase motors, they exhibit extensive diversity depending on the presence of an outer membrane, the structure of the cell wall, and the nature of their substrates. The FtsK/VirD4-type ATPase motor protein is a conserved component in the T4SS apparatus and forms a hexameric ring that allows substrate to pass through a central pore. Pore size varies by cognate substrate [54]. Consistent with other FtsK/ VirD4 homologs, TraD_pELF_ is predicted to form a homohexameric ring structure, through which the DNA substrate passes during conjugation (Fig 7A). Furthermore, modeling predicts that TraC_pELF_ also forms a stable hexamer with a central ring pore (Fig 7B). Protein docking simulation using ClusPro (https://cluspro.org) suggests that the TraC and TraD hexamers form a ring complex [55] (Fig 7C). Interestingly, the diameter of the ring pore in the TraC_pELF_/TraD_pELF_ complex is estimated to be closer to that of the FtsK/VirD4 ring in SLP2 and larger than that of the circular pIP501 plasmid (Supplementary Fig S7). This difference may reflect distinct substrate sizes; pIP501 transfers single-stranded DNA via an RCR-like mechanism, further supporting the idea that pELF-type plasmids, such as SLP2, are transferred via the first-end model [51] (Fig 7D).

Genetic manipulation is a powerful methodology to establish direct evidence of gene function. In *E faecalis*, sophisticated genetic engineering tools, which were established based on knowledge of pheromone-responsive conjugative plasmids, have allowed extensive investigation of plasmid and cellular functions in *E. faecalis* [56]. Meanwhile, despite the recent increase in its clinical importance, genetic engineering methodologies for *E. faecium* remain relatively underdeveloped [57]. In this study, we successfully developed and implemented effective genetic engineering tools for *E. faecium*, which, in addition to the techniques established to date, are expected to provide new options for genetic studies of *E. faecium* and thereby advance the field. [58,59]. The pELF-type linear plasmid family is one of the most common plasmid families, acting as a carrier of the critical antimicrobial resistance genes worldwide. The insights into plasmid biology established in this study will facilitate a better understanding of the plasmid-mediated adaptive evolution of *E. faecium* strains.

## Materials and Methods

### Bacterial strains, plasmids and antimicrobial agents

The bacterial strains, plasmids, and oligonucleotides used in this study are listed in Supplementary Tables S1, S2, and S3, respectively. Enterococcal strains were routinely grown in brain-heart infusion (BHI) broth (Difco, Detroit, MI, USA) at 37°C [60] or 28°C to maintain temperature-sensitive plasmids. Negative selection for plasmid-harboring strains *pheS\** was carried out on MM9YG agar containing 10 mM *p*-chlorophenylalanine. *Escherichia coli* strains were grown in Luria-Bertani (LB) medium (Difco) at 37°C or 28°C to maintain temperature-sensitive plasmids. Antibiotics used to select *E. coli* included 100 mg/L ampicillin (ABPC), 30 mg/L chloramphenicol (CP), and 20 mg/L gentamicin (GEN). The concentrations used for the routine selection of *E. faecium* harboring pMGJK72, pMGJK53, or pMGJK55 derivatives were 500 mg/L GEN. All antibiotics were obtained from Sigma-Aldrich (St. Louis, MO, USA).

### Total RNA isolation

An overnight bacterial culture was inoculated into BHI at a 10-fold dilution and pre-incubated at 37°C for 1 h. Subsequently, 500 µL of the preculture broth was inoculated into 4.5 mL of fresh BHI broth and incubated at 37°C for 3 or 8 h to reach exponential and stationary phases, respectively. For the co-culture conditions, the recipient strain was also preincubated and mixed at an equal volume. After incubation, an equal volume (5 mL) of RNAprotect (Qiagen) reagent was added to the culture broth and mixed by inversion. Stabilized cells were collected by centrifugation at 5,000 × *g* for 10 min. Total RNA was extracted from the pellet using the RNeasy Plus Kit (Qiagen) according to the manufacture’s instructions. The obtained RNA was analyzed using a Qubit fluorometer or a nanophotometer to determine concentration and purity, respectively.

### RNA sequencing and data analysis

Starting with 100 ng of total RNA, rRNA was depleted using the NEBNext rRNA Depletion Kit (Bacteria) (NEB, Ipswich, MA). Stranded cDNA libraries were prepared using the Illumina Stranded mRNA Prep Ligation kit (Illumina, San Diego, CA) according to the manufacture’s instructions and sequenced on a NovaSeq 6000 platform using the NovaSeq 6000 SP Reagent Kit v1.5 (300 cycles) (Illumina) with a 151 bp-paired-end protocol. The raw reads were analyzed using FastQC and mapped to the pELF2 sequence (<u>AP022343.1</u>) using Bowtie2 (v. 2.5.3) with default parameters.

Differential expression analysis was performed using DESeq2 (v.1.48.0). Data visualization was carried out using the ggplot2 package in R.

### Construction of the pMGJK72 plasmid and its derivatives for genetic manipulation

To construct a vector for improved site-directed genetic engineering in *E. faecium*, the following three DNA fragments were assembled. The first fragment containing the temperature-sensitive origin (*ori_pWO01_TS*), was PCR-amplified using primer pair (JK773/ JK804) and pGPA1 [61] as a template. The second fragment containing the gentamicin-resistance gene *aacA-aphD* for selection (*gen^r^*) was PCR-amplified using primer pair (JK774/JK747) and pGPA1 as a template. The third fragment containing the suicide gene (a mutant form of *pheS*) and a multiple cloning site (MCS) for Golden Gate Assembly (*pheS*\*+MCS) was PCR-amplified using primer pair (JK465/JK748) and pMGJK09 as a template. The first fragment was digested with *Bsa*I, whereas the second and the third fragments were digested with *BsmB*I. The fragments were ligated using T4 DNA ligase. The resulting plasmid after gentamicin selection in *E. coli* HST08 was confirmed by whole-plasmid sequencing and designated pMGJK72.

To construct pMGJK72 derivatives for site-directed mutagenesis of each *tra*_pELF_ gene, 1-kb DNA fragments flanking the upstream or downstream regions of each gene were PCR-amplified using the primer pairs JK857/JK858, JK934/JK860, JK898/JK899, JK900/JK901, JK902/JK903, JK904/JK905, JK906/JK907, JK908/JK909, JK914/JK915, JK916/JK917, JK918/JK919, JK920/JK921, JK926/JK927, and JK928/JK929 for *traC*, *traD*, *traE*, *traF*, *traH*, *traI*, and *traK*, respectively and the KUHS13 genomic DNA as a template. The resulting fragments and the pMGJK72 plasmid were digested with *Bsa*I and ligated using T4 DNA ligase. The resulting plasmids were selected on gentamicin in *E. coli* HST08 and confirmed via Sanger sequencing.

### Construction of the pMGJK53 plasmid and its derivatives for *in trans* IPTG-inducible expression

To construct an IPTG-inducible *in trans* expression, a linear DNA fragment was generated using the primer pair JK704/JK705 and the pRecT plasmid as a template. The obtained DNA fragment contains the whole pRecT plasmid [58] excluding the *recT* gene, was digested with *Bsa*I and dephosphorylated using Quick CIP (NEB). The super-folder *gfp* (*sfgfp*) gene was PCR-amplified using primer pair JK532/JK533 and a synthetic DNA fragment (Integrated DNA Technologies, Inc., Coralville, IA) as a template. The obtained *sfgfp* DNA fragment was digested with *BsmB*I and ligated to the pRecT-derived DNA fragment using T4 DNA ligase. The resulting plasmid after spectinomycin selection in *E. coli* HST08 was designated pMGJK51. Notably, the cloned *sfgfp* gene contained an internal *Bsa*I site that generates AGTC and GATT sticky ends for Golden Gate Assembly. The resulting plasmid after spectinomycin selection in *E. coli* HST08 was designated pMGJK51.

To promote the expression efficiency of the cloned genes via efficient transcriptional termination, pMGJK51 was linearized by PCR using primers JK533 and JK708, followed by digestion with *Bsa*I. The *rrnB* terminator sequence was amplified using primer pair JK530/JK531 and pTurbo as a template, followed by digestion with *BsmB*I. The *rrnB* terminator fragment was ligated into the linearized pMGJK51 using T4 DNA ligase. The resulting plasmid selected on spectinomycin in *E. coli* HST08 was designated pMGJK52.

To replace the spectinomycin selection marker with a gentamicin selection marker, pMGJK52 was linearized by PCR using primer pair JK711/JK712. The obtained DNA fragment, which contained the whole pMGJK52 plasmid except for the *spc^r^* gene, was digested with *Bsa*I. The fragment of gentamicin-resistance gene *aacA-aphD* for selection (*gen^r^*) was amplified using primer pair JK774/JK747 and pGPA1 as a template. The obtained *gen^r^* DNA fragment was digested with *Bsa*I and ligated into the pMGJK52-derived DNA fragment using T4 DNA ligase. The resulting plasmid after gentamicin selection in *E. coli* HST08 was confirmed via whole-plasmid sequencing and designated pMGJK53.

To construct pMGJK53 derivatives for ectopic expression of individual *tra*_pELF_ gene, each gene was PCR-amplified using combination of primer pairs JK993/JK994, JK995/JK996, JK885/JK886, JK997/JK998, JK999/JK1000, JK1001/JK1002, JK1003/JK1004, JK1005/JK1006, JK1007/JK1008, JK1009/JK1010, JK1011/JK1012, and JK1013/JK1014 for *traA*, *traB*, *traC*, *traD*, *traE*, *traF*, *traG*, *traH*, *traI, traJ*, *traK*, and *traL* respectively, using KUHS13 genomic DNA as a template. The resulting fragments and pMGJK53 plasmid were digested with *Bsa*I and ligated using T4 DNA ligase. The resulting plasmids after gentamicin selection in *E. coli* HST08 were confirmed via Sanger sequencing.

To construct pMGJK53 derivatives expressing mScarlet-I fusion with TraC or TraD, an *mScarlet-I* gene that is codon-optimized for *Lactococcus lactis* codon was synthesized (IDT Inc., IA) and PCR-amplified using a combination of JK602 and JK1025, followed by the addition of a linker sequence via PCR using primer pair JK1025/JK832. The resulting linker-*mScarlet-I* fragment was PCR-amplified using JK1025 and JK1026 to introduce *Bsa*I sites for ligation with the genes of interest. The *traC* and *traD* genes were PCR-amplified using primer pairs JK885/JK1029 and JK997/JK1030, respectively. These fragments were assembled via digestion with *Bsa*I followed by ligation using T4 DNA ligase, and subsequently cloned into pMGJK53 as described above. The resulting plasmids after gentamicin selection in *E. coli* HST08 were confirmed via Sanger sequencing.

### Construction of the pMGJK55 plasmid and its derivatives for promoter assay with luciferase reporter gene

To construct a promoter activity reporter plasmid based on pMGJK53, a synthetic luciferase gene, codon-optimized for *L. lactis*, was PCR-amplified using primer pair JK757/JK758 (IDT Inc.). The amplified fragment was then ligated into pMGJK53 via Golden Gate Assembly using *Bsa*I as described above. To remove the *lacI* repressor gene, pMGJK53::*luc* was linearized by PCR using JK735 and JK812, and then re-circularized via sequential treatment with *Bsa*I and T4 DNA ligase. The resulting plasmid pMGJK53::*luc*_Δ*lacI* was again linearized via PCR using primer pair JK745/JK746 and digested with *Bsa*I. Annealed oligonucleotides JK1015 and JK1016 were digested with *Bsa*I and mixed with the linearized pMGJK53::*luc*_Δ*lacI*, followed by ligation using T4 DNA ligase. The resulting plasmid after gentamicin selection in *E. coli* HST08 was confirmed via whole-plasmid sequencing and designated pMGJK55.

To construct pMGJK55 derivative harboring the *tra* promoter regions (P*tra*), the upstream region of *traA* with various lengths, P*tra*1, P*tra*2 or P*tra*3 was PCR-amplified using primers JK1022 and JK1019, JK10120 or JK1021, respectively, and KUHS13 genomic DNA as template.

### Site-directed recombination using pMGJK72 derivatives

The pMGJK72 derivatives were introduced into *E. faecium* KUHS13 via electroporation, followed by selection on BHI agar containing gentamicin (500 mg/L) at 30°C. Selected colonies obtained at the permissive temperature (30°C), where the plasmids are supposed to be maintained outside the host genome, were then streaked on pre-warmed BHI agar containing gentamicin (500 mg/L) and incubated at 42°C. Visible colonies obtained at the non-permissive temperature, where the plasmids are supposed to be integrated into the host genome, were collected and re-streaked at least twice in the same conditions to ensure the integration. The resulting colonies were inoculated into 5 mL BHI broth without any selection and incubated at 37°C overnight. One hundred microliters of a 100-fold dilution of the overnight culture was plated on MM9YG agar containing 10 mM *p*-chlorophenylalanine, followed by incubation at 37°C overnight. Colonies obtained from the negative selection, where the plasmids are supposed to be cured, were genotypically screened via PCR. The desired genetic deletions were finally confirmed using Sanger sequencing.

### Conjugative transfer

Conjugation assays were performed as described previously [60]. KUHS13 or its isogenic derivatives were used as the donor strains. *E. faecium* BM4105RF was used as the recipient strain. Briefly, donor and recipient strains were grown overnight in BHI at 37°C. The overnight culture was diluted 10-fold in fresh BHI and incubated for an additional 1 h at 37°C. A mixture was prepared by combining 250 μL each of donor and recipient cultures. For filter mating, 500 µL of the donor and recipient mixture was passed through a 0.22 μm nitrocellulose filter (Merck Millipore, Darmstadt, Germany) and placed onto BHI agar without antibiotic selection. After incubation at 37°C for 5 h, the bacterial cells on the nitrocellulose filter were collected in 1 mL phosphate-buffered saline (PBS) by vortexing. For broth mating, 500 µL of the donor and recipient mixture was incubated in a microcentrifuge tube without agitation at 37°C for 6 h. After incubation in either condition, serial dilution of the cell suspension was plated on a selective BHI agar containing rifampicin (25 mg/L), fusidic acid (25 mg/L), and VAN (16 mg/L) or BHI agar containing VAN (16 mg/L) to count the colonies of the transconjugant or donor, respectively, and then incubated at 37°C for 48 h. The colony count on the agar plate was determined, and the estimated ratio of the transconjugant CFU count to that of the donor was calculated as the conjugative transfer efficiency.

### Growth analysis and luciferase expression assay

To measure the kinetics of bacterial growth and reporter gene expression, an overnight culture of the *E. faecium* strain was inoculated into fresh BHI at a 100-fold dilution. To detect bioluminescence activity derived from the luciferase reporter gene expression, D-luciferin was added to the medium at a final concentration of 10 µg/ml. A 200 µL of the bacterial suspension was aliquoted into each well of a 96-well white microplate with a clear bottom, with three replicates per sample. Bioluminescence and optical density at 620 nm were monitored every 10 min during incubation at 37°C, using a heater-equipped plate reader Spark (Tecan, Switzerland).

### Fluorescence microscopic observation

Overnight culture of *E. faecium* strain was inoculated into fresh BHI containing 500 mg/L gentamicin and 0.1 mM IPTG, followed by incubation at 37°C for 2 h. The cells were then washed with PBS and chemically fixed using 4% paraformaldehyde. After fixation, the cells were mounted on 1% agar in PBS, layered onto a glass slide. The observations were performed using a fluorescence microscope (Axiovert 200; Carl Zeiss, Oberkochen, Germany), and the images were acquired using a DP71 camera (Olympus, Tokyo, Japan). The obtained images were processed and analyzed as previously described [62].

### Phylogenetic analysis

All FASTA-formatted nucleotide sequences of *E. faecium* (Taxonomy ID: 1352) deposited were retrieved from the RefSeq microbial genome database [43] using SeqKit (v.2.10.0) [63]. Of these sequences, contigs in which the 500-bp nucleotide sequence of the hairpin-end of the pELF2 was detected using BLASTn (v.2.16.0) [64], were extracted as pELF-like plasmid sequences. The CDSs in the extracted sequences were annotated via Prokka (v.1.14.6) [65]. Pangenomic analysis was performed via Roary (v.3.13.0) with a 70% identity parameter setting [66]. Phylogenetic tree accompanied by gene presence/absence panels was constructed using the ggtree package [67,68]. Synteny plot was generated using DiGAlign (https://www.genome.jp/digalign/) [69].

## Acknowledgments

pRecT_2 was a gift from Howard Hang (Addgene plasmid #167546; http://n2t.net/addgene:167546; RRID: Addgene_167546). pGPA1 was a gift from Willem van Schaik (Addgene plasmid #115476; http://n2t.net/addgene:115476; RRID: Addgene_115476).

## References

1. Eichel VM, Last K, Brühwasser C, Baum H von, Dettenkofer M, Götting T, et al. Epidemiology and outcomes of vancomycin-resistant enterococcus infections: a systematic review and meta-analysis. J Hosp Infect. 2023;141: 119–128. doi:10.1016/j.jhin.2023.09.008

2. Tacconelli E, Carrara E, Savoldi A, Harbarth S, Mendelson M, Monnet DL, et al. Discovery, research, and development of new antibiotics: the WHO priority list of antibiotic-resistant bacteria and tuberculosis. Lancet Infect Dis. 2018;18: 318–327. doi:10.1016/s1473-3099(17)30753-3

3. Almeida-Santos AC, Novais C, Peixe L, Freitas AR. Vancomycin-resistant Enterococcus faecium: A current perspective on resilience, adaptation, and the urgent need for novel strategies. J Glob Antimicrob Resist. 2025;41: 233–252. doi:10.1016/j.jgar.2025.01.016

4. Tomita H, Lu J-J, Ike Y. High Incidence of Multiple-Drug-Resistant Pheromone-Responsive Plasmids and Transmissions of VanA-Type Vancomycin-Resistant Enterococcus faecalis between Livestock and Humans in Taiwan. Antibiot (Basel, Switz). 2023;12: 1668. doi:10.3390/antibiotics12121668

5. Hashimoto Y, Hisatsune J, Suzuki M, Kurushima J, Nomura T, Hirakawa H, et al. Elucidation of host diversity of the VanD-carrying genomic islands in enterococci and anaerobes. Jac-antimicrobial Resist. 2022;4: dlab189. doi:10.1093/jacamr/dlab189

6. Hisatsune J, Tanimoto K, Kohara T, Myoken Y, Tomita Y, Sugai M. First Isolation of Vancomycin-Resistant Enterococcus faecium Carrying Plasmid-Borne vanD1. Antimicrob Agents Chemother. 2022;66: e01029–22. doi:10.1128/aac.01029-22

7. Hashimoto Y, Taniguchi M, Uesaka K, Nomura T, Hirakawa H, Tanimoto K, et al. Novel Multidrug-Resistant Enterococcal Mobile Linear Plasmid pELF1 Encoding vanA and vanM Gene Clusters From a Japanese Vancomycin-Resistant Enterococci Isolate. Front Microbiol. 2019;10: 2568. doi:10.3389/fmicb.2019.02568

8. Nomura T, Hashimoto Y, Kurushima J, Hirakawa H, Tanimoto K, Zheng B, et al. New colony multiplex PCR assays for the detection and discrimination of vancomycin-resistant enterococcal species. J Microbiol Methods. 2018;145: 69–72. doi:10.1016/j.mimet.2017.12.013

9. Zheng B, Tomita H, Inoue T, Ike Y. Isolation of VanB-type Enterococcus faecalis strains from nosocomial infections: first report of the isolation and identification of the pheromone-responsive plasmids pMG2200, encoding VanB-type vancomycin resistance and a Bac41-type bacteriocin, and pMG2201, encoding erythromycin resistance and cytolysin (Hly/Bac). Antimicrobial Agents and Chemotherapy. 2009;53: 735–747. doi:10.1128/aac.00754-08

10. Li Y, Tomita H, Lv Y, Liu J, Xue F, Zheng B, et al. Molecular characterization of erm(B)- and mef(E)-mediated erythromycin-resistant Streptococcus pneumoniae in China and complete DNA sequence of Tn2010. Journal of Applied Microbiology. 2010;110: 254–265. doi:10.1111/j.1365-2672.2010.04875.x

11. Tomita H, Ike Y. Genetic Analysis of the Enterococcus Vancomycin Resistance Conjugative Plasmid pHT : Identification of the Region Involved in Cell Aggregation and traB, a Key Regulator Gene for Plasmid Transfer and Cell Aggregation. Journal of bacteriology. 2008;190: 7739–7753. doi:10.1128/jb.00361-08

12. Zheng BB, Tomita H, Xiao YHY, Wang SS, Li YY, Ike Y. Molecular characterization of vancomycin-resistant Enterococcus faecium isolates from mainland China. Journal of Clinical Microbiology. 2007;45: 2813–2818. doi:10.1128/jcm.00457-07

13. Takeuchi KK, Tomita H, Fujimoto S, Kudo MM, Kuwano H, Ike Y. Drug resistance of Enterococcus faecium clinical isolates and the conjugative transfer of gentamicin and erythromycin resistance traits. FEMS microbiology letters. 2005;243: 8–8. doi:10.1016/j.femsle.2004.12.022

14. Tomita H, Ike Y. Genetic analysis of transfer-related regions of the vancomycin resistance Enterococcus conjugative plasmid pHTbeta: identification of oriT and a putative relaxase gene. Journal of bacteriology. 2005;187: 7727–7737. doi:10.1128/jb.187.22.7727-7737.2005

15. Tomita H, Tanimoto K, Hayakawa S, Morinaga K, Ezaki K, Oshima H, et al. Highly conjugative pMG1-like plasmids carrying Tn1546-like transposons that encode vancomycin resistance in Enterococcus faecium. J Bacteriol. 2003;185: 7024–8. doi:10.1128/jb.185.23.7024-7028.2003

16. Tomita H, Pierson C, Lim SK, Clewell DB, Ike Y. Possible connection between a widely disseminated conjugative gentamicin resistance (pMG1-like) plasmid and the emergence of vancomycin resistance in Enterococcus faecium. Journal of Clinical Microbiology. 2002;40: 3326–3333.

17. Ike Y, Tanimoto K, Tomita H, Takeuchi K, Fujimoto S. Efficient transfer of the pheromone-independent Enterococcus faecium plasmid pMG1 (Gmr) (65.1 kilobases) to Enterococcus strains during broth mating. J Bacteriol. 1998;180: 4886–92. doi:10.1128/jb.180.18.4886-4892.1998

18. Tanimoto K, Tomita H, Ike Y. The traA gene of the Enterococcus faecalis conjugative plasmid pPD1 encodes a negative regulator for the pheromone response. Plasmid. 1996;36: 55–61. doi:10.1006/plas.1996.0032

19. Fujimoto S, Tomita H, Wakamatsu E, Tanimoto K, Ike Y. Physical mapping of the conjugative bacteriocin plasmid pPD1 of Enterococcus faecalis and identification of the determinant related to the pheromone response. Journal of bacteriology. 1995;177: 5574.

20. Allen F, McInnes RS, Schaik W van, Moran RA. IS1216 drives the evolution of pRUM-like multidrug resistance plasmids in Enterococcus faecium. Microb Genom. 2025;11: 001598. doi:10.1099/mgen.0.001598

21. Segawa T, Masuda K, Hisatsune J, Ishida-Kuroki K, Sugawara Y, Kuwabara M, et al. Genomic analysis of inter-hospital transmission of vancomycin-resistant Enterococcus faecium sequence type 80 isolated during an outbreak in Hiroshima, Japan. Antimicrob Agents Chemother. 2024;68: e01716–23. doi:10.1128/aac.01716-23

22. Hisatsune J, Tanimoto K, Kohara T, Myoken Y, Tomita Y, Sugai M. First Isolation of Vancomycin-Resistant Enterococcus faecium Carrying Plasmid-Borne vanD1. Antimicrob Agents Chemother. 2022;66: e01029–22. doi:10.1128/aac.01029-22

23. Lebreton F, Schaik W van, McGuire AM, Godfrey P, Griggs A, Mazumdar V, et al. Emergence of epidemic multidrug-resistant Enterococcus faecium from animal and commensal strains. mBio. 2013;4: e00534-13-13-e00534-13. doi:10.1128/mbio.00534-13

24. Willems RJ, Schaik W van. Transition of Enterococcus faecium from commensal organism to nosocomial pathogen. Futur Microbiol. 2009;4: 1125–1135. doi:10.2217/fmb.09.82

25. Daza MVB, Cortimiglia C, Bassi D, Cocconcelli PS. Genome-based studies indicate that the Enterococcus faecium Clade B strains belong to Enterococcus lactis species and lack of the hospital infection associated markers. Int J Syst Evol Microbiol. 2021;71. doi:10.1099/ijsem.0.004948

26. Mathpal S, Panickar A, Joshi T, Ramaiah S, Anbarasu A. Genomic surveillance of vancomycin-resistant Enterococcus faecium: a study on Resistome, Plasmidome, and mobilome profiling. Curr Genet. 2025;71: 26. doi:10.1007/s00294-025-01331-y

27. Kurushima J, Nomura T, Ota N, Tomita H. Complete genomes of clade A1 and B Enterococcus faecium isolates harboring pHTβ, a vanA-type vancomycin-resistant pMG1-like plasmid. Microbiol Resour Announc. 2025;14: e00684–25. doi:10.1128/mra.00684-25

28. Zaidi S-Z, Zaheer R, Poulin-Laprade D, Scott A, Rehman MA, Diarra M, et al. Comparative Genomic Analysis of Enterococci across Sectors of the One Health Continuum. Microorganisms. 2023;11: 727. doi:10.3390/microorganisms11030727

29. Hashimoto Y, Suzuki M, Kobayashi S, Hirahara Y, Kurushima J, Hirakawa H, et al. Enterococcal Linear Plasmids Adapt to Enterococcus faecium and Spread within Multidrug-Resistant Clades. Antimicrob agents Chemother. 2023;67: e0161922. doi:10.1128/aac.01619-22

30. Almeida-Santos AC, Tedim AP, Duarte B, Silva LM, Teixeira J, Castro AP, et al. Unnoticed spread of linear VanA-plasmids in vancomycin-variable Enterococcus faecium strains across different regions: a diagnostics challenge. J Antimicrob Chemother. 2025; dkaf409. doi:10.1093/jac/dkaf409

31. Hashimoto Y, Dao DT, Kasuga I, Takemura T, Abe H, Hasebe F, et al. Ongoing independent evolution of linezolid and vancomycin-resistance pELF-type linear plasmids across the One Health spectrum. Antimicrob Agents Chemother. 2025;69: e01168–25. doi:10.1128/aac.01168-25

32. Kent AG, Spicer LM, Campbell D, Breaker E, McAllister GA, Ewing TO, et al. Sentinel Surveillance reveals phylogenetic diversity and detection of linear plasmids harboring vanA and optrA among enterococci collected in the United States. Antimicrob Agents Chemother. 2024;68: e00591–24. doi:10.1128/aac.00591-24

33. Bakthavatchalam YD, Puraswani M, Livingston A, Priya M, Venkatesan D, Sharma D, et al. Novel linear plasmids carrying vanA cluster drives the spread of vancomycin resistance in Enterococcus faecium in India. J Glob Antimicrob Resist. 2022;29: 168–172. doi:10.1016/j.jgar.2022.03.013

34. Cinthi M, Coccitto SN, Simoni S, Gherardi G, Palamara AT, Lodovico SD, et al. The optrA, cfr(D) and vanA genes are co-located on linear plasmids in linezolid- and vancomycin-resistant enterococcal clinical isolates in Italy. J Antimicrob Chemother. 2025;80: 1362–1370. doi:10.1093/jac/dkaf082

35. Beh JQ, Daniel DS, Judd LM, Wick RR, Kelley P, Cronin KM, et al. Genomics to understand the global landscape of linezolid resistance in Enterococcus faecium and Enterococcus faecalis. Microb Genom. 2025;11: 001432. doi:10.1099/mgen.0.001432

36. Fujiya Y, Harada T, Sugawara Y, Akeda Y, Yasuda M, Masumi A, et al. Transmission dynamics of a linear vanA-plasmid during a nosocomial multiclonal outbreak of vancomycin-resistant enterococci in a non-endemic area, Japan. Sci Rep. 2021;11: 14780. doi:10.1038/s41598-021-94213-5

37. Sun L, Zhuang H, Chen M, Chen Y, Chen Y, Shi K, et al. Vancomycin heteroresistance caused by unstable tandem amplifications of the vanM gene cluster on linear conjugative plasmids in a clinical Enterococcus faecium. Antimicrob Agents Chemother. 2024;68: e01159–23. doi:10.1128/aac.01159-23

38. Boumasmoud M, Haunreiter VD, Schweizer TA, Meyer L, Chakrakodi B, Schreiber PW, et al. Genomic Surveillance of Vancomycin-Resistant Enterococcus faecium Reveals Spread of a Linear Plasmid Conferring a Nutrient Utilization Advantage. mBio. 2022;13: e03771–21. doi:10.1128/mbio.03771-21

39. Grohmann E, Christie PJ, Waksman G, Backert S. Type IV secretion in Gram-negative and Gram-positive bacteria. Mol Microbiol. 2018;107: 455–471. doi:10.1111/mmi.13896

40. Chen CW, Yu T, Lin Y, Kieser HM, Hopwood DA. The conjugative plasmid SLP2 of Streptomyces lividans is a 50 kb linear molecule. Mol Microbiol. 1993;7: 925–932. doi:10.1111/j.1365-2958.1993.tb01183.x

41. Hsu C-C, Chen CW. Linear Plasmid SLP2 Is Maintained by Partitioning, Intrahyphal Spread, and Conjugal Transfer in Streptomyces. J Bacteriol. 2009;192: 307–315. doi:10.1128/jb.01192-09

42. Hashimoto Y, Kita I, Suzuki M, Hirakawa H, Ohtaki H, Tomita H. First Report of the Local Spread of Vancomycin-Resistant Enterococci Ascribed to the Interspecies Transmission of a vanA Gene Cluster-Carrying Linear Plasmid. mSphere. 2020;5. doi:10.1128/msphere.00102-20

43. Tatusova T, Ciufo S, Fedorov B, O’Neill K, Tolstoy I. RefSeq microbial genomes database: new representation and annotation strategy. Nucleic Acids Res. 2014;42: D553–D559. doi:10.1093/nar/gkt1274

44. Weaver KE. Enterococcal Genetics. Microbiol Spectr. 2019;7. doi:10.1128/microbiolspec.gpp3-0055-2018

45. Sterling AJ, Snelling WJ, Naughton PJ, Ternan NG, Dooley JSG. Competent but complex communication: The phenomena of pheromone-responsive plasmids. Plos Pathog. 2020;16: e1008310. doi:10.1371/journal.ppat.1008310

46. Breuer RJ, Hirt H, Dunny GM. Mechanistic Features of the Enterococcal pCF10 Sex Pheromone Response and the Biology of Enterococcus faecalis in Its Natural Habitat. J Bacteriol. 2018;200: 10.1128/jb.00733-17. doi:10.1128/jb.00733-17

47. Tanimoto K, Ike Y. Analysis of the Conjugal Transfer System of the Pheromone-Independent Highly Transferable Enterococcus Plasmid pMG1: Identification of a tra Gene (traA) Up-Regulated during Conjugation. J Bacteriol. 2002;184: 5800–5804. doi:10.1128/jb.184.20.5800-5804.2002

48. Fraikin N, Couturier A, Lesterlin C. The winding journey of conjugative plasmids toward a novel host cell. Curr Opin Microbiol. 2024;78: 102449. doi:10.1016/j.mib.2024.102449

49. 49. Garcillán - Barcia MP, Francia MV, Cruz FDL. The diversity of conjugative relaxases and its application in plasmid classification. FEMS Microbiol Rev. 2009;33: 657–687. doi:10.1111/j.1574-6976.2009.00168.x

50. Xu M, Zhu Y, Zhang R, Shen M, Jiang W, Zhao G, et al. Characterization of the Genetic Components of Streptomyces lividans Linear Plasmid SLP2 for Replication in Circular and Linear Modes. J Bacteriol. 2006;188: 6851–6857. doi:10.1128/jb.00873-06

51. Lee H-H, Hsu C-C, Lin Y-L, Chen CW. Linear plasmids mobilize linear but not circular chromosomes in Streptomyces: support for the ‘end first’ model of conjugal transfer. Microbiology. 2011;157: 2556–2568. doi:10.1099/mic.0.051441-0

52. Jumper J, Evans R, Pritzel A, Green T, Figurnov M, Ronneberger O, et al. Highly accurate protein structure prediction with AlphaFold. Nature. 2021;596: 583–589. doi:10.1038/s41586-021-03819-2

53. Costa TRD, Patkowski JB, Macé K, Christie PJ, Waksman G. Structural and functional diversity of type IV secretion systems. Nat Rev Microbiol. 2024;22: 170–185. doi:10.1038/s41579-023-00974-3

54. Paillard P, Rouger Q, Thomet M, Macé K. Type IV secretion systems: from structures to mechanisms. EMBO J. 2025;44: 6304–6319. doi:10.1038/s44318-025-00584-0

55. Kozakov D, Hall DR, Xia B, Porter KA, Padhorny D, Yueh C, et al. The ClusPro web server for protein–protein docking. Nat Protoc. 2017;12: 255–278. doi:10.1038/nprot.2016.169

56. Kristich CJ, Chandler JR, Dunny GM. Development of a host-genotype-independent counterselectable marker and a high-frequency conjugative delivery system and their use in genetic analysis of Enterococcus faecalis. Plasmid. 2007;57: 131–144. doi:10.1016/j.plasmid.2006.08.003

57. Kurushima J, Tomita H. Advances of genetic engineering in streptococci and enterococci. Microbiol Immunol. 2022;66: 411–417. doi:10.1111/1348-0421.13015

58. Chen V, Griffin ME, Maguin P, Varble A, Hang HC. RecT Recombinase Expression Enables Efficient Gene Editing in Enterococcus spp. Appl Environ Microbiol. 2021;87: e00844–21. doi:10.1128/aem.00844-21

59. de Maat V, Stege PB, Dedden M, Hamer M, van Pijkeren J-P, Willems RJL, et al. CRISPR-Cas9-mediated genome editing in vancomycin-resistant Enterococcus faecium. Fems Microbiol Lett. 2019;366. doi:10.1093/femsle/fnz256

60. Dunny GM, Craig RAR, Carron RLR, Clewell DB. Plasmid transfer in Streptococcus faecalis: production of multiple sex pheromones by recipients. Plasmid. 1979;2: 454–465.

61. Zhang X, Maat V de, Prieto AMG, Prajsnar TK, Bayjanov JR, Been M de, et al. RNA-seq and Tn-seq reveal fitness determinants of vancomycin-resistant Enterococcus faecium during growth in human serum. BMC Genom. 2017;18: 893. doi:10.1186/s12864-017-4299-9

62. Schindelin J, Arganda-Carreras I, Frise E, Kaynig V, Longair M, Pietzsch T, et al. Fiji: an open-source platform for biological-image analysis. Nature methods. 2012;9: 676–682. doi:10.1038/nmeth.2019

63. Shen W, Sipos B, Zhao L. SeqKit2: A Swiss army knife for sequence and alignment processing. iMeta. 2024;3: e191. doi:10.1002/imt2.191

64. Camacho C, Coulouris G, Avagyan V, Ma N, Papadopoulos J, Bealer K, et al. BLAST+: architecture and applications. BMC Bioinform. 2009;10: 421. doi:10.1186/1471-2105-10-421

65. Seemann T. Prokka: rapid prokaryotic genome annotation. Bioinformatics. 2014;30: 2068–2069. doi:10.1093/bioinformatics/btu153

66. Page AJ, Cummins CA, Hunt M, Wong VK, Reuter S, Holden MTG, et al. Roary: rapid large-scale prokaryote pan genome analysis. Bioinformatics. 2015;31: 3691–3693. doi:10.1093/bioinformatics/btv421

67. Yu G, Smith DK, Zhu H, Guan Y, Lam TT. ggtree: an r package for visualization and annotation of phylogenetic trees with their covariates and other associated data. Methods Ecol Evol. 2017;8: 28–36. doi:10.1111/2041-210x.12628

68. Wickham H. ggplot2, Elegant Graphics for Data Analysis. 2016. doi:10.1007/978-3-319-24277-4

69. Nishimura Y, Yamada K, Okazaki Y, Ogata H. DiGAlign: Versatile and Interactive Visualization of Sequence Alignment for Comparative Genomics. Microbes Environ. 2024;39: ME23061. doi:10.1264/jsme2.me23061

